# Oxytocin enhancement of emotional empathy: generalization across cultures and effects on amygdala activity

**DOI:** 10.1101/307256

**Authors:** YaYuan Geng, Weihua Zhao, Feng Zhou, Xiaole Ma, Shuxia Yao, Rene Hurlemann, Benjamin Becker, Keith M. Kendrick

## Abstract

Accumulating evidence suggests that the neuropeptide oxytocin can enhance empathy although it is unclear which specific behavioral and neural aspects are influenced, and whether the effects are modulated by culture, sex and trait autism. Based on previous findings in Caucasian men, we hypothesized that a single intranasal dose of oxytocin would specifically enhance emotional empathy via modulatory effects on the amygdala in an Asian (Chinese) population and explored the modulatory role of sex and trait autism on the effects. We first conducted a double-blind, randomized between-subject design experiment using a modified version of the multifaceted empathy task (MET) to determine whether oxytocin’s facilitation of emotional empathy can be replicated in Chinese men (n = 60). To further explore neural mechanisms behind and potential sex differences, functional MRI and skin conductance measures were acquired in an independent experiment incorporating men and women (n = 72). Oxytocin enhanced emotional empathy across experiments and sex, an effect that was accompanied by reduced amygdala activity and increased skin conductance responses. On the network level oxytocin enhanced functional coupling of the right amygdala with the insula and posterior cingulate cortex for positive valence stimuli but attenuated coupling for negative valence stimuli. The effect of oxytocin on amygdala functional connectivity with the insula was modulated by trait autism. Overall, our findings provide further support for the role of oxytocin in facilitating emotional empathy and demonstrate that effects are independent of culture and sex and involve modulatory effects on the amygdala and its interactions with other key empathy regions.

## 1. Introduction

Empathy is a key social-cognitive capacity that facilitates interpersonal functioning by allowing us to recognize, understand and respond appropriately to mental and affective states experienced by others (Decety and Jackson, 2004; Dziobek, et al., 2008; Reniers, et al., 2010). Impaired empathy is a core deficit in psychiatric disorders characterized by interpersonal dysfunctions, including autism (Dziobek, et al., 2008), schizophrenia (Lee, et al., 2011; Rosenfeld, et al., 2011; Shamay-Tsoory, et al., 2007), and personality disorders (Herpertz and Bertsch, 2014).

Empathy is a multidimensional construct, entailing cognitive processes of perspective-taking, to make inferences about others’ mental states (cognitive empathy, CE), as well as emotional processes reflecting a direct affective reaction involving understanding, sharing and responding appropriately to others’ feelings (emotional empathy, EE) (Bernhardt and Singer, 2012; Shamay-Tsoory, 2011; Shamay-Tsoory, et al., 2009). EE has been further divided into a direct component (direct emotional empathy, EED), referring to explicit emotional evaluation and empathic concern, and an indirect component (indirect emotional empathy, EEI), referring to a more general physiological arousal response to both person and context (Dziobek, et al., 2008). Although the cognitive and emotional components of empathy represent partly dissociable systems (Shamay-Tsoory, et al., 2009), integrative approaches propose that the experience of empathy evolves as a dynamic interplay between them requiring an explicit representation of the specific affective state of the other person, thereby making CE a prerequisite for EE (Decety and Jackson, 2004; Hillis, 2014). On the neural level the functional organization of empathy is partially mirrored in shared and separable anatomical representations (Bernhardt and Singer, 2012; Lamm, et al., 2011; Lamm, et al., 2007; Leigh, et al., 2013; Schulte-Ruther, et al., 2007; Singer and Lamm, 2009), with the bilateral insula, posterior cingulate cortex (PCC) and anterior cingulate cortex (ACC) contributing to both (Fan, et al., 2011), and the amygdala contributing to the emotional component of empathy (Cox, et al., 2012; Leigh, et al., 2013).

Converging evidence suggests that the hypothalamic neuropeptide oxytocin (OXT) facilitates empathy (Riem, 2012; Rosenfeld, et al., 2011; Striepens, et al., 2011). Genetic approaches have consistently revealed associations between individual variations in the OXT receptor gene and levels of trait empathy in Caucasian (Rodrigues, et al., 2009; Smith, et al., 2014) and Chinese populations (Wu, et al., 2012), with more recent studies suggesting that the specific associations evolve in interaction with other factors, particularly culture (Luo, et al., 2015b; Montag, et al., 2017) and sex (Weisman, et al., 2015). Studies investigating the behavioral effects of intranasal OXT administration on CE have reported enhanced accuracy in the reading the mind in the eyes test (RMET) (Domes, et al., 2007b) and a paradigm requiring participants to infer the intensity of positive or negative emotions expressed by subjects portrayed in videos (Bartz, et al., 2010). However, findings in the domain of CE have been variable, with OXT effects in the RMET being either restricted to difficult items (Feeser et al., 2015) or unable to be reproduced at all even when taking into account stimulus difficulty and valence (Radke and de Bruijn, 2015). Other studies have also reported that effects were more pronounced in individuals with poor baseline performance (Riem, et al., 2014) or high trait autism (Bartz, et al., 2010). Studies that aimed specifically at determining effects of OXT on EE focused on empathy for pain, an evolutionary conserved primary emotional component (Decety, 2011; Panksepp and Panksepp, 2013), and found no effect on pain empathy towards a partner (Singer, et al., 2008), although an enhanced pain empathic response towards members of an out-group (Shamay-Tsoory, et al., 2013). In contrast, another study in men using the Multifaceted Empathy Test (MET) (Dziobek, et al., 2008), which assesses both CE and EE, observed that OXT specifically enhanced both EED and EEI, but not CE (Hurlemann, et al., 2010). This latter study additionally demonstrated selective EE deficits in amygdala lesion patients and therefore suggested that the amygdala may mediate the EE enhancing effects of OXT. Although several neuroimaging studies have demonstrated modulatory effects of intranasally administered OXT on the core neural components of the empathy network, including the insula, ACC and amygdala and their functional interactions, across different task paradigms (Bakermans-Kranenburg and van IJzendoorn, 2013; Herpertz and Bertsch, 2016; Wigton, et al., 2015), to date only two studies have directly explored the neural mechanisms underlying OXT’s empathy enhancing effects. The first reported that OXT increased activation in the superior temporal gyrus and insula during the RMET task (Riem, et al., 2014), whereas the second reported reductions in left insula activity during pain empathic processing (Bos, et al., 2015).

In summary, although the empathy enhancing effects of OXT are central to its proposed social-cognitive and therapeutic properties, it remains unclear whether it selectively enhances CE or EE, and which specific neural substrates are involved. To systematically address these questions, we employed two independent pharmacological between-subject placebo (PLC) controlled experiments in healthy Chinese individuals investigating the effects of intranasal OXT on CE and EE and the underlying neural basis of this effects during the MET (Dziobek, et al., 2008).

Previous studies on the empathy enhancing potential of intranasal OXT are entirely based on observations in Caucasian populations. However, there is accumulating evidence from OXT-administration studies either employing comparable experimental protocols in Caucasian and Chinese subjects (Hurlemann, et al., 2010; Hu, et al., 2015) or examining moderating effects of key cultural orientation differences such as a collectivistic orientation (Pfundmair, et al., 2014; Xu, et al., 2017), suggesting culture-dependent social-cognitive effects of OXT. To this end, the first experiment aimed to replicate findings in a male Caucasian sample showing that OXT enhances EE but not CE (Hurlemann, et al., 2010) in a male Chinese sample. In a second independent sample, male and female Chinese participants performed the same MET paradigm during fMRI to determine the neural substrates involved. Analyses on the neural level focused on the insula, amygdala and ACC as core empathy regions (Bernhardt and Singer, 2012; Lamm, et al., 2011; Lamm, et al., 2007; Leigh, et al., 2013; Schulte-Ruther, et al., 2007; Singer and Lamm, 2009). Given that the amygdala has been specifically (Cox, et al., 2012; Leigh, et al., 2013) and critically (Hurlemann, et al., 2010) associated with emotional facets of empathy, we expected that OXT’s enhancement of EE would be accompanied by altered regional activity and network level connectivity of the amygdala. Previous studies reported increased as well as decreased amygdala activity and connectivity following OXT (Domes, et al., 2007a; Hu, et al., 2015; Striepens, et al., 2012; Tully, et al., 2018; Wigton, et al., 2015) therefore no directed hypothesis with respect to OXT’s neural effect was formulated.

Based on a growing number of findings suggesting sex-dependent effects of OXT on social cognition (Gao, et al., 2016; Chen, et al., 2016; Luo, et al., 2017), the second experiment additionally explored whether OXT differentially affects empathic processing in men and women. In line with a previous study reporting that sex does not affect OXT’s modulation of empathy (Shamay-Tsoory, et al., 2013), we hypothesized that OXT facilitation of EE would generalize across sexes. Finally, in the context of increasing interest in the therapeutic application of OXT as a potential treatment to improve social cognitive deficits, including empathy, in autism spectrum disorders (Young and Barrett, 2015), and in line with previous studies in healthy subjects (Bartz, et al., 2010; Scheele, et al., 2014; Xu, et al., 2015), the modulatory role of trait autism (assessed by the Autism Spectrum Quotient questionnaire, ASQ, Baron-Cohen, et al., 2001) was explored.

## 2. Materials and Methods

### 2.1 Participants

To fully replicate the previous study on Caucasian participants (Hurlemann, et al., 2010), only males were recruited in the first experiment but both males and females were enrolled in the second experiment to explore potential sex-dependent effects of OXT on empathy. Experiment 1 (Exp 1) included 60 participants (M ± SD, mean age = 22.42 ± 2.23, all male) and Experiment 2 (Exp 2) included an independent sample of 72 participants (34 females, mean age = 21.18 ± 1.95, 38 males, mean age = 22.61 ± 2.01). Both experiments incorporated a double-blind, between-participant design, with participants being randomly assigned to receive either OXT or PLC nasal-spray, resulting in n = 30 (Exp 1) and n = 36 (Exp 2, female = 17) participants treated with OXT. The experimental groups in both experiments were of comparable age (Exp 1, p = 0.53, T_58_ = 0.63; Exp 2, p = 0.66, T_70_ = - 0.44), education (Exp 1, p = 0.66, T_58_ = 0.44; Exp 2, p = 0.63, T_70_ = -0.49) and, in Exp 2, of equivalent sex distribution (chi-square < 0.001, df = 1, p = 1). Exclusion criteria for all participants were past or current physical, neurological or psychiatric disorders, regular or current use of medication or tobacco.

Participants were required to abstain from alcohol, caffeine or nicotine for at least 12 hours before the experiment. None of the females in Exp 2 were taking oral contraceptives or were tested during their menstrual period. Menstrual cycle phase was determined using validated procedures as described in (Penton-Voak, et al., 1999). The proportion of females estimated to be in their follicular or luteal phases did not differ significantly between the treatment groups (chi-square = 0.12, df = 1, p = 0.73). In Exp 1, one participant (in the OXT group) and in Exp 2, three participants (in the PLC group) failed to understand task instructions and were consequently excluded from all further analysis, leading to a total of n = 59 participants in Exp 1 and n = 69 participants in Exp 2.

Before the experiment, written informed consent was obtained from all participants. The study was approved by the local ethics committee of the University of Electronic Science and Technology of China and all procedures were in accordance with the latest revision of the declaration of Helsinki.

### 2.2 Experimental Protocol

To control for potential confounding variables, all participants initially completed the following questionnaires: Becks Depression Inventory (BDI; Beck, et al., 1961), WLEIS-C Emotional Intelligence Scale (Wleis-C; Wong and Law, 2002), State Trait Anxiety Inventory (STAI; Spielberger, et al., 1970), Empathy Quotient (EQ; Baron-Cohen and Wheelwright, 2004), and Positive and Negative Affect Scale (PANAS; Watson, et al., 1988). To examine associations with trait autism, the ASQ questionnaire (Baron-Cohen, et al., 2001) was administered. Intranasal treatment (oxytocin nasal spray, Sichuan Meike Pharmacy Co., Ltd., China, or placebo nasal spray with identical ingredients except oxytocin) was administered in line with recommendations for the intranasal administration of OXT in humans (Guastella, et al., 2013) and 45min before the start of the experimental paradigm. In Exp 1, three puffs per nostril (at 30s intervals) were administered (24 IU) and in Exp 2, five puffs per nostril (40 IU). Both doses are in the typical range employed by other studies (Guastella, et al., 2013; Striepens, et al., 2011) with the rationale for increasing the dose in Exp 2 being to explore dose-dependent behavioral effects of OXT. In a previous study we found equivalent behavioral and neural effects of 24 and 48 IU OXT doses (Zhao, et al., 2017). However, it should be noted that findings from some other studies investigating dose-dependent effects of intranasal OXT have suggested an inverted-U-shaped dose-response curve (Cardoso, et al., 2013; Quintana, et al., 2017; Quintana, et al., 2016; Spengler, et al., 2017) and thus a stronger enhancement of EE with 24 IU relative to 40 IU is conceivable. In post experiment interviews, participants were unable to guess better than chance whether they had received the OXT nasal spray, confirming successful blinding.

### 2.3 Experimental paradigm

In line with a previous study on male Caucasian participants (Hurlemann, et al., 2010), empathy was assessed using the MET (Dziobek et al., 2008; Domes, et al., 2013; Edele, et al., 2013; Hurlemann, et al., 2010; Wingenfeld, et al., 2014), which assesses both EE and CE components using ecologically valid photo-based stimuli of either negative or positive valence. To account for potential confounding effects of OXT on in-group versus out-group empathy (De Dreu and Kret, 2016) and a cultural empathy bias (Cao, et al., 2015; Luo, et al., 2015a), the original Caucasian MET stimuli were exchanged with corresponding pictures displaying Chinese protagonists. The Chinese stimuli were initially evaluated in an independent sample (supplementary materials) and the final set of Chinese stimuli (30 positive, 30 negative valence) closely resembled the Caucasian stimuli depicting daily life scenarios and conveying emotional mental states via facial expression, body posture and contextual cues. To assess CE, participants were instructed to infer the emotional state of the protagonist in each scene and choose the corresponding answer from 4 options listed. The 4 options presented similar but distinct emotional states to ensure at least 70% accuracy for each stimulus picture. For EED, participants were required to rate how they felt for the protagonist in the depicted scene (1-9 scale, 1 = not at all, 9 = very strong), for EEI participants were required to rate how much they were aroused by the scene (1-9 scale, 1 = very calm, 9 = very aroused).

The different components of empathy were presented in a mixed event/block-design. Following a 3 second(s) instruction cue and a jittered inter-trial interval of 3.9s (2.3-5.9s), 10 stimuli per block were each presented for 3s followed by either a choice of the emotion depicted for the CE condition (displayed for 4 s) or a rating scale (1-9) for the EED and EEI conditions (displayed for 5 s). Six blocks were presented for each condition, resulting in a total of 18 blocks. The order of blocks was counterbalanced across the experimental conditions, and the fMRI experiment was divided into 6 runs, each containing one block per empathy component. During fMRI (Exp 2) electrodermal activity was simultaneously acquired as an index of autonomic sympathetic activity (Stern, et al., 2001) (technical details on the electrodermal data acquisition are provided in the supplementary materials). To allow baseline recovery of the electrodermal signal a mean inter-trial interval of 5s (4-6s) and a mean interval separating stimulus presentation and behavioral response of 4s (3-5s) was adopted for the fMRI experiment.

### 2.4 fMRI data acquisition

The fMRI data in Exp 2 were collected using a GE (General Electric Medical System, Milwaukee, WI, USA) 3.0T Discovery 750 MRI scanner. fMRI time series were acquired using a T2*-weighted echo planar imaging pulse sequence (repetition time, 2000 millisecond (ms); echo time, 30ms; slices, 39; thickness, 3.4 millimeter (mm); gap, 0.6 mm; field of view, 240 × 240 mm^2^; resolution, 64×64; flip angle, 90°). Additionally, a high resolution T1-weighted structural image was acquired using a 3D spoiled gradient recalled (SPGR) sequence (repetition time, 6 ms; echo time, 2ms; flip angle 9°; field of view, 256 × 256 mm^2^; acquisition matrix, 256 × 256; thickness, 1 mm without gap) to exclude participants with apparent brain pathologies and to improve normalization of the fMRI data.

### 2.5 fMRI data processing

fMRI data were analyzed using SPM12 (Wellcome Trust Center of Neuroimaging, University College London, London, United Kingdom). The first five volumes were discarded to allow T1 equilibration and images were realigned to the first image to correct for head motion. Tissue segmentation, bias-correction and skull-stripping were done for the high-resolution structural images. The functional time series were co-registered with the skull-stripped anatomical scan and normalized to MNI space with voxel size of 3 mm^3^. Normalized images were then spatially smoothed using a Gaussian kernel with full-width at half-maximum (FWHM) of 8 mm. On the first level, event-related responses were modelled and subsequently convolved with the standard hemodynamic response function (HRF). The first level design matrix included valence-(positive, negative) and empathy type-(CE, EED, EEI) specific regressors for the viewing phases as main experimental conditions. In addition, regressors for the cue presentation, valence-and empathy type-specific regressors for the rating phases, and for viewing and rating phases of incorrect trials as well as the six movement regressors were included. The experimental contrasts were next submitted to a second level random effects analysis.

To evaluate empathy-type specific main and interaction effects of treatment and valence, repeated-measured ANOVAs were employed in a flexible-factorial design. Based on our regional hypothesis and the core empathy network (Bakermans-Kranenburg and van IJzendoorn, 2013; Bernhardt and Singer, 2012; Hillis, 2014; Hurlemann, et al., 2010; Leigh, et al., 2013; Shamay-Tsoory and Abu-Akel, 2016; Wigton, et al., 2015), the analyses focused on the bilateral amygdala, insula and ACC which were structurally defined using 60% probability maps from the Harvard-Oxford (sub)cortical atlas. For the regionally focused analysis approach condition-specific parameter estimates were extracted from these regions of interest (ROI) using the Marsbar toolbox (Brett, et al., 2002) and subjected to empathy type-specific ANOVAs with the between-participant factor treatment (OXT, PLC) and the within-participant factor valence (positive, negative) in SPSS (Statistical Package for the Social Sciences, Version 22). P-values for the post-hoc tests of the ROI analysis were Bonferroni-corrected (P < 0.05). An exploratory voxel-wise whole-brain analysis in SPM that served to determine contributions of brain regions outside of the predefined network of interest was thresholded at P < 0.05, corrected using the family-wise error (FWE) approach.

To investigate the effects of OXT on the network level, a generalized form of psychophysiological interaction analysis (gPPI; http://brainmap.wisc.edu/PPI; McLaren, et al., 2012) was conducted using regions showing significant OXT effects in the BOLD level analysis as seeds and implementing an empathy-type specific voxel-wise whole-brain ANOVA approach including the between-participant factor treatment (OXT, PLC) and the within-participant factor valence (positive, negative) thresholded at P < 0.05, FWE-corrected at the cluster level. In line with recent recommendations for the control of false-positives in cluster-based correction approaches an initial cluster forming threshold of P < 0.001 was applied to data with a resolution of 3 × 3 × 3 mm (Eklund, et al., 2016; Slotnick, 2017). Parameter estimates were extracted from the significant regions to disentangle the specific effects in post-hoc comparisons. Finally, associations between neural indices and trait autism (ASQ scores) were conducted in SPSS using Pearson correlation analysis.

## 3. Results

In both experiments, there were no significant differences in trait and mood questionnaire scores between OXT and PLC treatment groups (supplementary Table S1 for Exp 1, Table S2 for Exp 2). In line with previous studies in Chinese populations (Melchers et al., 2015; Montag et al., 2017), no significant sex differences in ASQ and EQ scores were observed in Exp 2 (Supplementary Table S3).

### 3.1 Behavioral results

Based on previous conceptualizations of empathy, proposing that CE is a prerequisite for EE (Decety and Jackson, 2004; Hillis, 2014), for EED and EEI measures only trials for which subjects successfully recognized the emotions displayed by the protagonist were analyzed (for a similar approach see (Luo, et al., 2015a)). To this end, correctly recognized trials were initially determined based on the CE performance, with only correct trials subsequently entering the analyses for the EED and EEI facets.

There were no significant differences in CE accuracy between the two treatment groups in both experiments (Exp 1, OXT, 77.53% ± 6.02%, PLC, 78.94% ± 5.78%, T_57_ = 0.92, p = 0.36; Exp 2, OXT, 80.10% ± 6.18%, PLC, 81.57% ± 5.83%, T_67_ = 1.02, p = 0.31). In Exp 1, there was a main effect of treatment (F (1, 57) = 6.46, p = 0.01, η^2^_p_ = 0.10) for EED indicating that OXT generally enhanced EED (Fig. 1A). There was no significant treatment × valence interaction (F (1, 57) = 0.96, p = 0.33, η^2^_p_ = 0.02). Analysis of EEI did not reveal a treatment main effect (F (1, 57) = 2.19, p = 0.14, η^2^_p_ = 0.04) or valence × treatment interaction effect (F (1, 57) = 3.16, p = 0.08, η^2^p = 0.05).

**Fig. 1.**
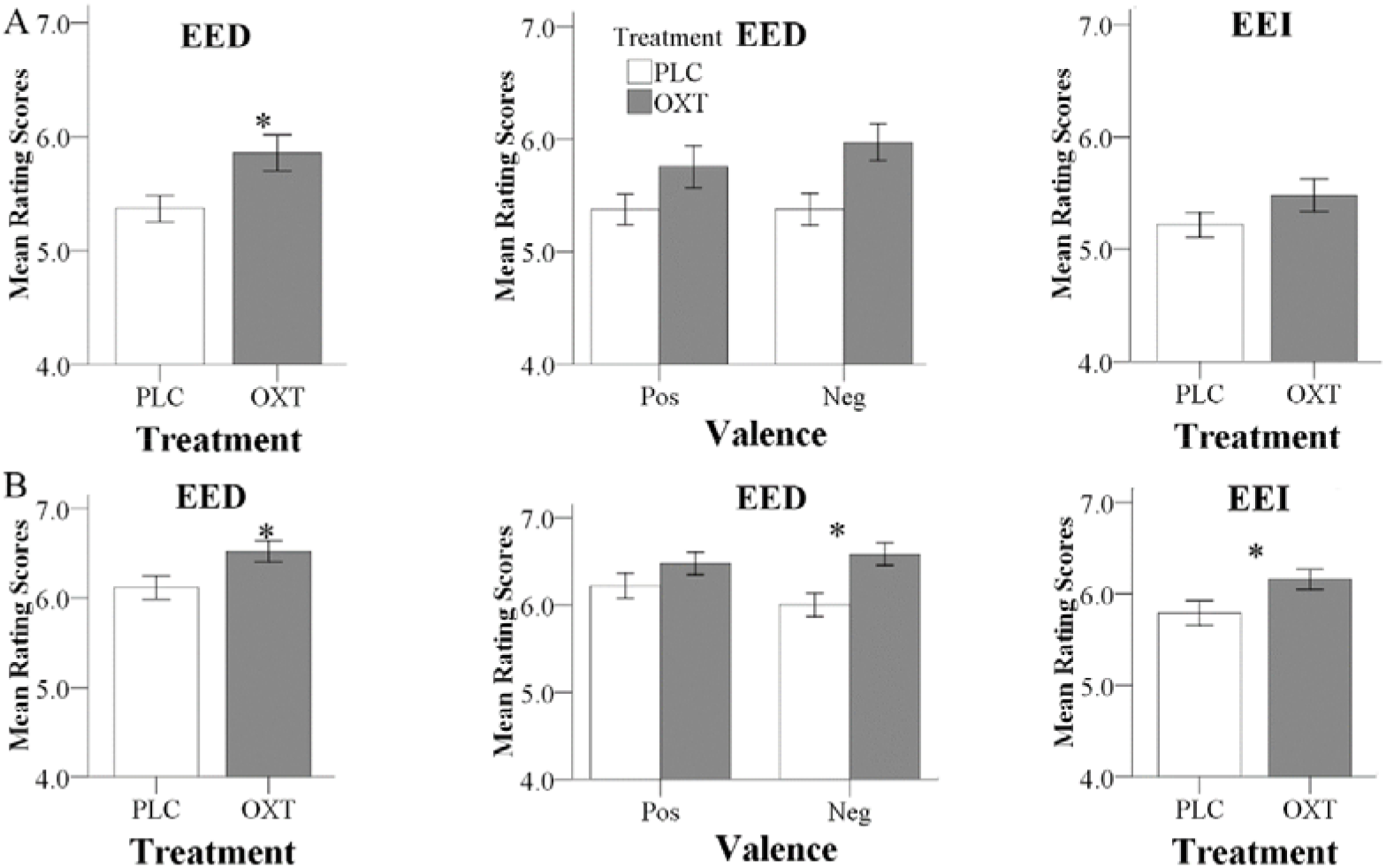
Behavioral results for Exp 1 (A) and Exp 2 (B). **(A)** In Exp 1, OXT significantly increased EED ratings. To allow for a better comparison with Exp 2, effects of OXT on positive and negative EED trials, as well as on EEI are also shown; **(B)** In Exp 2, OXT increased both EED and EEI ratings; an effect of oxytocin on negative EED drove the significant treatment by valence interaction. (*P < 0.05)

Consistent with the findings for EED in Exp 1, Exp 2 also yielded a significant main effect of treatment on EED (F (1, 67) = 5.81, p = 0.02, η^2^_p_ = 0.08) with higher ratings following OXT compared to PLC (Fig. 1B). There was also a significant valence × treatment interaction (F (1, 67) = 4.18, p = 0.05, η^2^_p_ = 0.06) with more pronounced effects of OXT on negative compared to positive valence stimuli (positive: F (1,67) = 1.68, p = 0.2, η^2^p = 0.02; negative: F (1, 67) = 9.96, p = 0.002, η^2^_p_ = 0.13). For EEI there was also a significant main effect of treatment (F (1, 67) = 4.84, p = 0.03, η^2^_p_ = 0.07) but no treatment × valence interaction (F (1, 67) = 2.02, p = 0.16, η^2^_p_ = 0.03). For CE, there were neither significant main effects nor interactions (F (1,65) = 0.80, p = 0.37, η^2^_p_ = 0.01). In Exp 2, no significant main or interaction effects involving sex were observed (all ps > 0.18) arguing against sex-dependent effects of OXT on empathy.

### 3.2 Associations between behavior and trait autism

In Exp 1, there was a trend towards a negative correlation between the ASQ score and the total EED and EEI scores in the OXT group (ASQ Total: EED r = -0.47, p = 0.09; EEI r = - 0.37, p = 0.19) but not the PLC group (EED r = 0.319, p = 0.18; EEI r = 0.17, p = 0.48). The correlation significantly differed between the PLC and OXT groups for EED (Fisher’s z = - 2.13, p = 0.03) although not for EEI (Fisher’s z = -1.43, p = 0.15). In Exp 2, there was a similar pattern of correlation differences between EED and EEI scores and total ASQ scores, although these associations did not reach statistical significance other than for EED under OXT (EED - PLC r = 0.03, p = 0.85, OXT r = -0.34, p = 0.04, Fisher’s z = 1.53, p = 0.13; EEI - PLC r = 0.03, p = 0.85; OXT r = -0.28, p = 0.09, Fisher’s z =1.29, p = 0.20).

Regression plots are shown in Fig. 2 and suggest that OXT is producing its main behavioral effects in participants with lower autism traits.

**Fig. 2.**
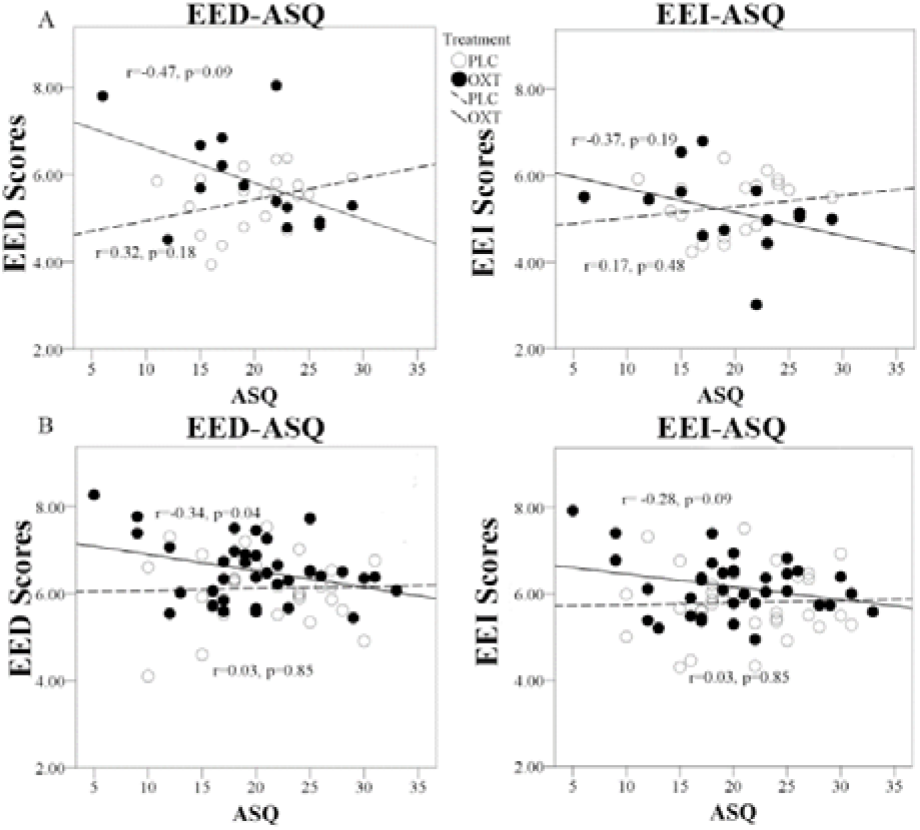
Regression plots for ASQ score and EED and EEI ratings. Exp 1 **(A)** and Exp 2 **(B)** in OXT and PLC groups.

### 3.3 Dose-dependent effects between Experiments 1 and 2 (24 IU vs 40 IU)

Dose effects were explored by combining the data from male participants in Exp 1 (24 IU) and Exp 2 (40 IU). To initially explore potential effects of the different experimental environments (Exp 1, 24 IU, behavioral testing room; Exp 2, 40 IU, inside the MRI-scanner) on empathy per se, a first analysis focused on the placebo-treated subjects. A repeated ANOVA with environment (behavioral vs MRI room) as a between-subject factor and valence as a within-subject factor revealed a significant environment main effect for both EE facets, indicating elevated EE ratings in the MRI room (Main effects: EED: p = 0.003, F (1,47) = 10.19, η^2^_p_ = 0.18; EEI: p = 0.02, F (1,47) = 5.74, η^2^_p_ = 0.11, both interactions with valence > 0.38, non-significant, Fig. 3), but no effects on CE (Main effect: p = 0.15, F (1,47) = 2.19, iļ^2^p = 0.05; Interaction: p = 0.9, F (1,47) = 0.02, η^2^p < 0.001). These findings suggest that the MRI-environment per se increased EE, an effect possibly related to elevated levels of stress during the MRI assessments, which would be in line with a previous study reporting that stress-induction specifically increased EE, but not CE in the MET (Wolf, et al., 2015). The environmental differences and potential interactions with OXT preclude the interpretation of dose-related differences between the experiments.

**Fig. 3.**
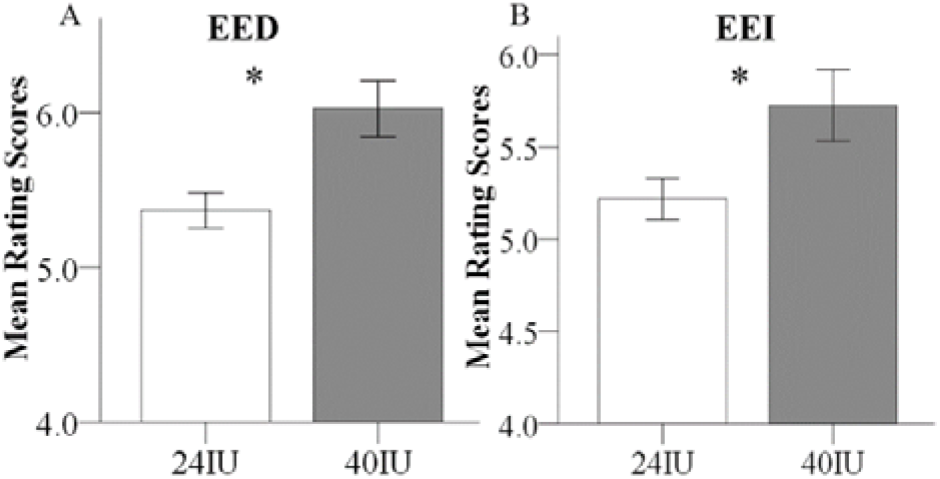
EE differences between Exp 1 and Exp 2 in the placebo treated subjects. Examining the placebo treated male subjects from the two experiments revealed that EE was significantly increased in the MRI environment (Exp 2, 40 IU) compared to the behavioral testing room (Exp 1, 24 IU).

### 3.4 Oxytocin effects on SCR

One participant was excluded from SCR analysis due to low skin impedance and thus a total of 68 participants from Exp 2 were included. Analyses of the SCR data paralleled the analyses of the empathy ratings, using ANOVAs with the between-subject factors treatment (OXT, PLC) and sex (male, female), and the within-subject factor valence. There was a marginal main effect of treatment on SCR during EED trials and significant during EEI trials (EED: F (1, 66) = 3.77, p = 0.06, η^2^p = 0.05; EEI: F (1, 66) = 4.50, p = 0.04, ifp = 0.06), but not during CE trials (F (1, 66) = 2.14, p = 0.15, η^2^p = 0.03). This was due to SCR responses being increased in the OXT group during EE and EEI trials (Fig. 4). There were no significant main effects of valence or treatment × valence or treatment × sex interactions for CE, EE or EEI trials (all ps>0.2).

**Fig. 4.**
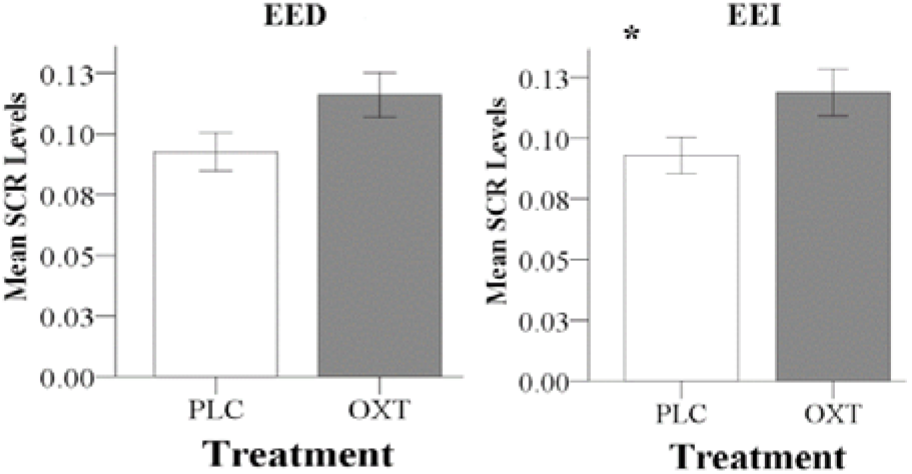
OXT effect on SCR in Exp 2. In Exp 2, OXT increased the SCR during both EED (P = 0.06) and EEI trials. (* P < 0.05)

### 3.5 Oxytocin effects on neural activity

In view of the absence of sex-dependent effects in the behavioral analysis, and to increase the statistical power to determine OXT effects on the neural level, the data from male and female participants were pooled for the fMRI analyses. Four further participants were excluded from the fMRI analysis due to excessive head motion (head motion > 3 mm). The neural mechanisms underlying the behavioral effects of OXT were initially explored in the different priori ROIs (amygdala, insula and ACC) using separate repeated measures ANOVAs for the three empathy (CE, EE and EEI) conditions. Main treatment effects were only observed in the amygdala (Fig. 5A) for EED (left amygdala: F (1,63) = 6.55, p = 0.01, η^2^p = 0.09; right amygdala: F (1,63) = 5.18, p = 0.03, η^2^_p_ = 0.08). There were no significant main effects for CE or EEI or any treatment × valence interactions for CE, EED or EEI. An exploratory whole brain analysis revealed no regions that showed significant treatment-dependent changes under CE, EED or EEI (all P_FDR___corrected_ > 0.05) outside of the prior defined ROIs.

**Fig. 5.**
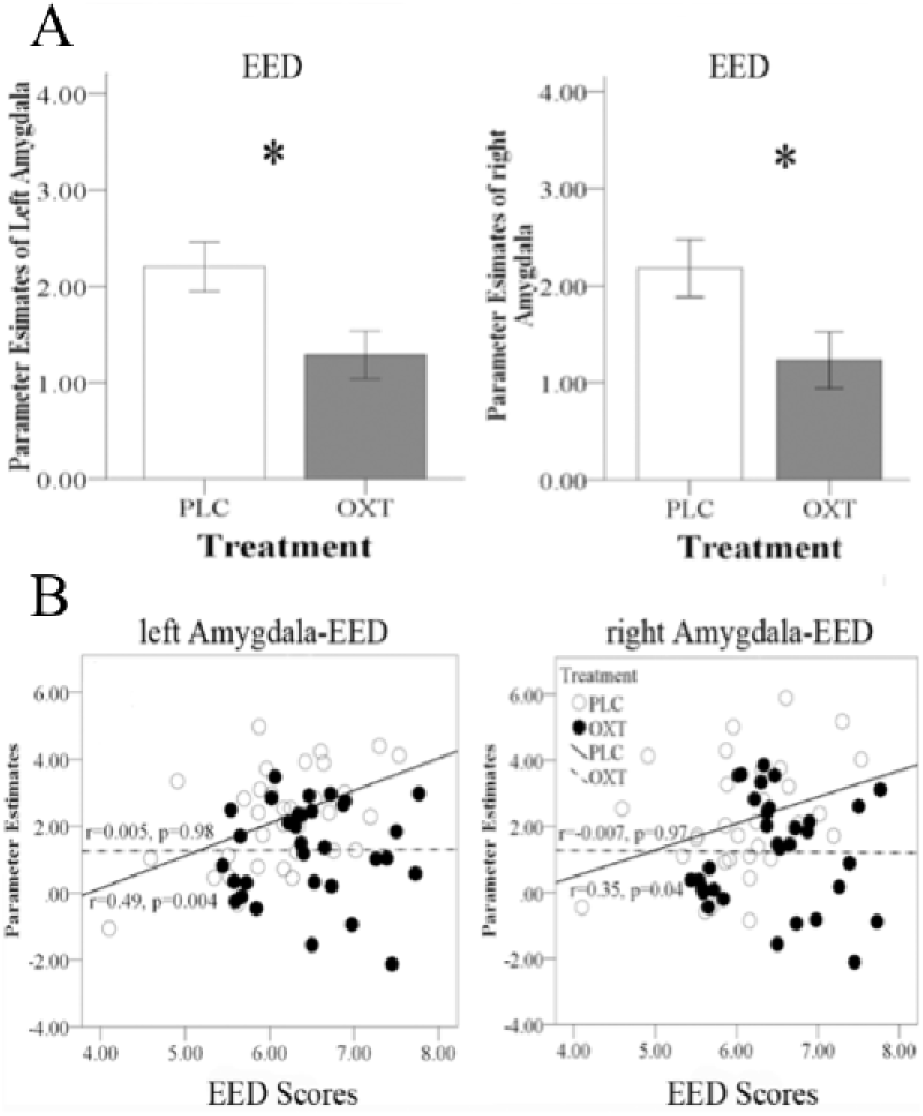
Effects of OXT on bilateral amygdala responses. **(A)** Region of interest analysis results for left and right amygdala responses during EED trials; **(B)** Regression plots show correlations between left and right amygdala responses and EED scores in OXT and PLC groups. *P < 0.05

### 3.6 Oxytocin effects on functional connectivity

Repeated measures ANOVA models in SPM that included the between-subject factors treatment (OXT, PLC) and sex (male, female) and the within-subject factor valence (positive, negative) revealed a significant Treatment × Valence interaction effect for EED-associated functional coupling of the right amygdala with the bilateral insula and the bilateral PCC (left insula peak located at x/y/z, -33/6/-15, P_FWE_ = 0.02, cluster size = 143 voxels; right insula peak located at 45/18/-12, P_FWE_ = 0.03, cluster size = 121 voxels; left PCC peak located at - 30/-33/30, P_FWE_ = 0.003, cluster size = 213 voxels; right PCC peak located at 21/-36/33, P_FWE_ = 0.01, cluster size = 149 voxels; coordinates given in MNI-space). Extraction of parameter estimates further revealed that OXT increased functional connectivity for positive valence stimuli whereas it decreased connectivity for negative valence ones (left insula: positive, F (1,61) = 10.52, p = 0.002, ifp = 0.15, negative, F (1,61) = 3.86, p = 0.05, η^2^p = 0.06; right insula: positive, F (1,61) = 3.34, p = 0.07, η^2^_p_ = 0.05, negative, F (1,61) = 5.53, p = 0.02, η^2^_p_ = 0.08; left PCC: positive, F (1,61) = 4.34, p = 0.04, η^2^p = 0.07, negative, F (1,61) = 5.67, p = 0.02, η^2^p = 0.09; right PCC: positive, F (1,61) = 8.00, p = 0.006, η^2^p = 0.12, negative, F (1,61) = 5.48, p = 0.02, η^2^p = 0.08) (Fig. 6).

**Fig. 6.**
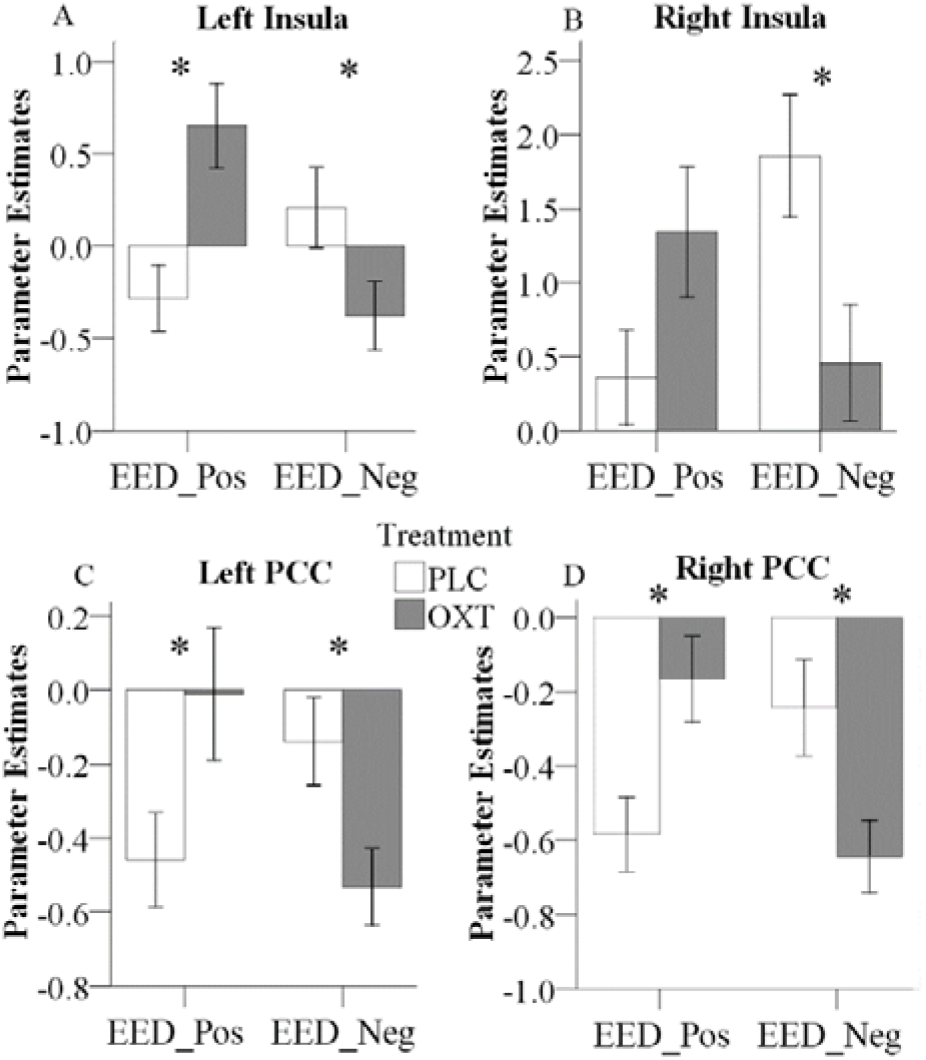
Effects of OXT effect on right amygdala functional connectivity. Effects of treatment on functional connectivity of the right amygdala, indicating valence-dependent effects of OXT on the coupling of the right amygdala with the left **(A)** and right **(B)** insula, as well as the left **(C)** and right **(D)** PCC (*P < 0.05). (Pos: Positive; Neg: Negative).

### 3.7 Associations between neural and SCR effects of OXT and behavioral and autism trait scores

There was a significant positive correlation between EED scores and bilateral amygdala responses in the PLC group (left r = 0.49, p = 0.004; right r = 0.35, p = 0.04) which was absent in the OXT group (left r = 0.005, p = 0.98; right r = -0.007, p = 0.97). The correlation difference between the PLC and OXT groups was significant for the left (Fisher’s Z = 2.05, p = 0.04) but not the right (Fisher’s Z = 1.43, p = 0.15) amygdala (Fig. 5B). There was no correlation between left or right amygdala responses with total ASQ scores.

For the functional connections showing OXT effects for EED in terms of a treatment × valence interaction, coupling strength between the right amygdala and left insula during positive valence EED trials was positively correlated with the total ASQ in the PLC group (total ASQ - r = 0.40, p = 0.02,) but not in the OXT group (total ASQ - r = -0.22, p = 0.22, Fisher’s Z = 2.50, p = 0.01; Fig. 7A). OXT particularly appears to increase the strength of right amygdala functional connections with the insula in individuals with lower ASQ scores, although only for positive valence EED. The strength of link between the right amygdala and left PCC during negative valence EED trials was positively correlated with the total ASQ score in the PLC group but not the OXT, although the difference between the groups was not significant (total ASQ - PLC r = 0.36, p = 0.04; OXT r = 0.15, p = 0.41; Fisher’s Z = 0.85, p = 0.39; Fig. 7B). There were no significant correlations between SCR values and ASQ scores during either EED or EEI trials (all ps > 0.41).

**Fig. 7.**
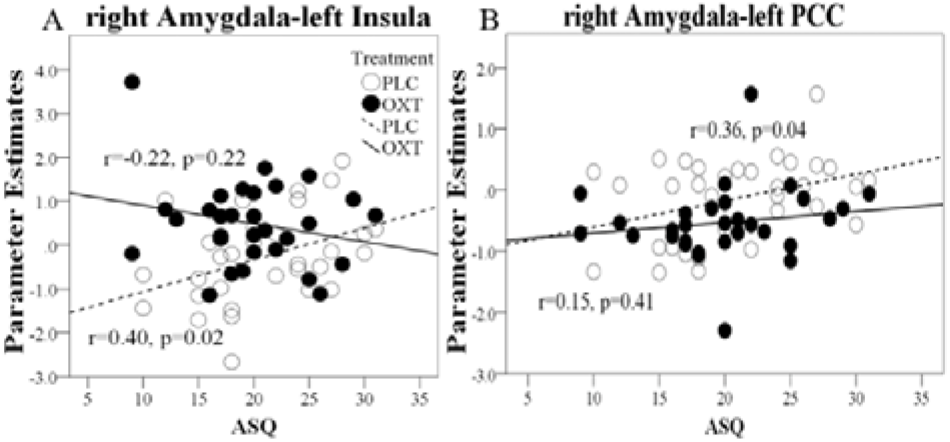
Regression plots for correlations of ASQ scores with amygdala responses and functional connectivity during EE trials in OXT and PLC groups. **(A)** Correlation between right amygdala-left PCC functional connectivity during negative EED trials and total ASQ score; **(B)** Correlation of right amygdala functional connectivity with left insula cortex during positive EE trials and total ASQ score. In all cases, significant positive correlations during PLC administration are absent in the OXT group.

## 4. Discussion

The present study confirmed in two independent samples that intranasal OXT specifically facilitates EE but not CE as assessed by the MET paradigm in Chinese participants, thereby replicating previous findings in Caucasian participants (Hurlemann, et al., 2010). Our findings also demonstrated for the first time that the OXT-induced enhancement of EED is associated with decreased bilateral amygdala reactivity and enhanced functional coupling of the right amygdala with the insula and PCC for positive valence stimuli but attenuated coupling for negative valence stimuli. These behavioral and neural effects were not modulated by subject sex, suggesting a generalization across men and women. Finally, an exploratory analysis of associations with trait autism revealed that both behavioral and neural effects of OXT were modulated to some extent by trait autism scores.

Although many studies have reported cultural differences between Asian and Caucasian participants in the context of OXT receptor polymorphisms and empathy (Jessica, et al., 2016; Kim, et al., 2010; Luo, et al., 2015b), we did not find any substantive difference with respect to the effects of intranasal OXT on empathy processing as assessed by MET. Thus, in both cultures, OXT enhanced EE but not CE (Hurlemann, et al., 2010) for both valences, although in our second experiment we found stronger effects for negative valence stimuli. Effects of the scanning environment on emotional empathy ratings per se precluded the direct evaluation of dose-response effects between the two experiments, however OXT specifically increased EE in both suggesting that its effects generalize across 24 and 40IU doses, in line with our previous finding (Zhao, et al., 2017). In general, the magnitude of the reported behavioral OXT effect on both EED and EEI reported in Caucasian participants was however somewhat stronger compared to both 24 and 40 IU doses administered in our study, although different MET stimuli were used.

In agreement with other studies, there were no sex-differences in EE, trait empathy (Wu et al., 2012) or trait autism (Kawamura et al., 2011; Montag et al., 2017) scores in our Chinese study cohort, whereas in Caucasian participants we found that females scored significantly higher than males for both positive and negative valence stimuli (Hurlemann, et al., 2010). Thus, it is conceivable that in Caucasian females the effects of OXT in the MET might not be as pronounced as in males. The absence of an effect of OXT on CE in the MET contrasts with reports using other paradigms, notably the reading the mind in the eyes test (RMET) (Domes, et al., 2007b; Feeser, et al., 2015). However, the robustness of these findings has been questioned by another study which failed to replicate them even when taking into account both item difficulty and valence (Radke and de Bruijn, 2015). Moreover, there are also notable differences between the MET and RMET with the images in the MET including more complex natural scenes and emotions conveyed by multiple cues (face, body posture and context) whereas in the RMET emotions are only interpreted from pictures of eye regions and are also often more subtle. Thus, OXT can facilitate CE in some contexts, particularly with cues restricted to eyes, but not in others where multiple cues are present. Additionally, and in contrast to previous studies, we measured SCR responses during trials involving the three empathy components and OXT only increased the SCR in EE and not CE trials. Thus, OXT enhancement of EE is paralleled by increased physiological arousal not only in EEI trials (where participants are asked to score how aroused they are by the stimulus picture) but also in EED trials (where they are scoring the strength of their feelings towards to protagonist in the picture).

In line with the specific, and critical contribution of the amygdala to emotional, rather than cognitive aspects of empathy (Hurlemann, et al., 2010), OXT’s enhancement of EE was accompanied by a reduction of associated amygdala activity. Exploratory analyses revealed that EED scores were positively associated with the magnitude of amygdala responses during positive valence trials in the PLC group, whereas this association was absent under OXT, possibly reflecting an enhancement of amygdala processing efficiency. While some previous studies found that OXT specifically reduced amygdala responses to negative emotional stimuli (Gamer, et al., 2010; Kirsch, et al., 2005), the suppression of EED-associated amygdala activity was observed irrespective of valence. A similar pattern of OXT-induced valence-independent suppression of amygdala activity has previously been suggested to reflect reduced uncertainty of a social stimulus which in turn motivates approach behavior (Domes, et al., 2007a). In line with this interpretation, the valence-independent EED-associated amygdala suppression may reflect that OXT’s approach-facilitating properties (Arakawa, et al., 2010) promote EE regardless of whether the emotions expressed by the protagonist are positive or negative, which is also in line with a rodent study reporting an overall reduction of amygdala EEG power following OXT (Sobota, et al., 2015). Other studies have found that OXT’s modulation of amygdala responses dependent upon sex (Gao, et al., 2016, Luo, et al., 2017) and it is generally considered that the salience of cues as well as their context may play an important role in determining OXT’s effects (Shamay-Tsoory and Abu-Akel, 2016). In the present study, neither sex nor valence influenced amygdala reactivity. This possibly reflects the fact that both salience and context are broadly similar for EE responses in the two sexes.

OXT also differentially altered the functional connectivity between the right amygdala and bilateral insula in a valence-dependent manner. In EED trials, the strength of the functional connectivity between the right amygdala and insula following OXT was significantly increased during positive valence stimuli but decreased during negative ones. A few previous studies have also reported OXT effects on functional connectivity between the insula and amygdala (Gao, et al., 2016; Hu, et al., 2015; Rilling, et al., 2012; Striepens, et al., 2012) and these two regions are key hubs of the brain salience network (Uddin, 2015). Thus, in the current context OXT may have acted to increase the salience of both positive and negative valence stimuli during EED trials by differentially altering the functional connectivity between the amygdala and insula. Rilling et al., (2012) have also previously suggested that the stronger the functional coupling between amygdala and insula, the more able the amygdala is to elicit subjective feeling states in response to salient social stimuli.

The effect of OXT on increasing functional connectivity between the right amygdala and bilateral PCC for positive valence stimuli and decreasing it for negative ones in EED trials may similarly reflect a modulatory influence on salience processing. A previous study has reported that OXT enhanced functional connectivity between amygdala and PCC during exposure to infant laughter (Riem, et al., 2012), suggesting that it increased the incentive salience of infant laughter. In our current study, the consistent patterns of functional connectivity changes elicited by OXT for positive and negative valence stimuli for amygdala functional connectivity with the insula and PCC may indicate that these three regions comprise an integrated network mediating valence-dependent OXT effects.

Both the behavioral and neural effects of OXT were modified to some extent by trait autism scores, as measured by the ASQ. In both experiments OXT tended to produce a negative correlation between EE and ASQ scores, whereas this correlation was absent in the PLC group. However, this effect of OXT only achieved significance in Exp 1, which included only male participants, and indicates that increased EE scores were more evident in individuals with lower ASQ scores. For the neural associations functional connectivity between the amygdala and insula was positively associated with total ASQ scores for positive valence EE trials in the PLC group, but this was absent in the OXT group. This indicates that OXT effects on functional connections between the right amygdala and left insula (for positive valence stimuli) were also strongest in individuals with lower ASQ scores. Thus overall, while both behavioral and neural OXT effects on EE were modified by ASQ scores, the extent to which these findings represent support for possible therapeutic use in ASD remains unclear. Indeed, a recent study on OXT enhancement of behavioral and neural responses to affective touch also reported stronger effects in individuals with lower ASQ scores (Scheele, et al., 2014).

There are several limitations which should be acknowledged in the current study. Firstly, we were unable to directly compare behavioral and neural responses during the MET task in Caucasian as well Chinese participants, so we cannot totally exclude the possibility that some cultural differences in response to OXT during empathic processing may exist. Secondly, we only investigated effects using the MET paradigm and it is possible that OXT effects on CE as well as EE would have been found using other paradigms. Lastly, the absence of sex-differences in OXT effects in the current study might have been contributed to by our Chinese male and female participants exhibiting similar EE scores, in contrast to Caucasian participants (Hurlemann, et al., 2010), and also similar ASQ scores.

In summary, in the current study we have shown that in the MET paradigm, OXT enhances EE but not CE in Chinese participants, similar to Caucasian ones, and additionally that this occurs in female as well as male participants. Furthermore, we have shown for the first time that this EE effect of OXT is associated with decreased amygdala responses and differentially altered functional connectivity between the amygdala and insula and PCC for positive and negative valence stimuli. Finally, we have shown that both behavioral and neural effects of OXT are modified to some extent by trait autism scores, although behavioral and functional connectivity effects were strongest in individuals with lower scores.

## Acknowledgements

This study was supported by the National Natural Science Foundation of China (NSFC) grant (grant number 31530032).

## Author Contributions Statement

Geng, Hurlemann and Kendrick designed this experiment, Geng collected the data, Geng, Zhao, Zhou, Kendrick and Becker analysed the data, Geng, Zhao, Ma, Yao, Becker and Kendrick interpreted the results. Geng, Becker and Kendrick wrote the paper.

## References

Arakawa, H., Arakawa, K., and Deak, T. (2010). Oxytocin and vasopressin in the medial amygdala differentially modulate approach and avoidance behavior toward illness-related social odor. Neuroscience 171(4), 1141–1151. doi: 10.1016/j.neuroscience.2010.10.013.

Bakermans-Kranenburg, M.J., and van IJzendoorn, M.H. (2013). Sniffing around oxytocin: review and meta-analyses of trials in healthy and clinical groups with implications for pharmacotherapy. Translational Psychiatry 3.

Baron-Cohen, S., and Wheelwright, S. (2004). The Empathy Quotient: An Investigation of Adults with Asperger Syndrome or High Functioning Autism, and Normal Sex Differences. Journal of Autism and Developmental Disorders 34(2), 163–175. doi: 10.1023/b:jadd. 0000022607.19833.00.

Baron-Cohen, S., Wheelwright, S., Skinner, R., Martin, J., and Clubley, E. (2001). The autism-spectrum quotient (AQ): evidence from Asperger syndrome/high-functioning autism, males and females, scientists and mathematicians. J Autism Dev Disord 31(1), 5–17.

Bartz, J.A., Zaki, J., Bolger, N., Hollander, E., Ludwig, N.N., Kolevzon, A., et al. (2010). Oxytocin Selectively Improves Empathic Accuracy. Psychological Science 21(10), 1426–1428. doi: 10.1177/0956797610383439.

Beck, A.T., Ward, C.H., Mendelson, M., Mock, J., and Erbaugh, J. (1961). An inventory for measuring depression. Arch Gen Psychiatry 4, 561–571.

Bernhardt, B.C., and Singer, T. (2012). The Neural Basis of Empathy. Annual Review of Neuroscience, Vol 35 35, 1–23.

Bethlehem, R.A.I., van Honk, J., Auyeung, B., and Baron-Cohen, S. (2013). Oxytocin, brain physiology, and functional connectivity: A review of intranasal oxytocin fMRI studies. Psychoneuroendocrinology 38(7), 962–974.

Bos, P.A., Montoya, E.R., Hermans, E.J., Keysers, C., and van Honk, J. (2015). Oxytocin reduces neural activity in the pain circuitry when seeing pain in others. Neuroimage 113, 217–224.

Brett, M., Anton, J.-L., Valabregue, R., and Poline, J.-B. (2002). Region of interest analysis using the MarsBar toolbox for SPM 99. Neuroimage 16(2), S497.

Cao, Y., Contreras-Huerta, L.S., McFadyen, J., and Cunnington, R. (2015). Racial bias in neural response to others' pain is reduced with other-race contact. Cortex 70, 68–78.

Cardoso, C., Ellenbogen, M.A., Orlando, M.A., Bacon, S.L., and Joober, R. (2013). Intranasal oxytocin attenuates the cortisol response to physical stress: a dose-response study. Psychoneuroendocrinology 38(3), 399–407. doi: 10.1016/j.psyneuen.2012.07.013.

Chen, X., Hackett, P.D., DeMarco, A.C., Feng, C., Stair, S., Haroon, E., et al. (2016). Effects of oxytocin and vasopressin on the neural response to unreciprocated cooperation within brain regions involved in stress and anxiety in men and women. Brain Imaging Behav 10(2), 581–593. doi: 10.1007/s11682-015-9411-7.

Cox, C.L., Uddin, L.Q., Di Martino, A., Castellanos, F.X., Milham, M.P., and Kelly, C. (2012). The balance between feeling and knowing: affective and cognitive empathy are reflected in the brain's intrinsic functional dynamics. Social Cognitive and Affective Neuroscience 7(6), 727–737. doi: 10.1093/scan/nsr051.

De Dreu, C.K., and Kret, M.E. (2016). Oxytocin Conditions Intergroup Relations Through Upregulated In-Group Empathy, Cooperation, Conformity, and Defense. Biol Psychiatry 79(3), 165–173. doi: 10.1016/j.biopsych.2015.03.020.

Decety J. (2011). The neuroevolution of empathy. Ann N Y Acad Sci 1231, 35–45. doi: 10.1111/j.1749-6632.2011.06027.x.

Decety, J., and Jackson, P.L. (2004). The functional architecture of human empathy. Behav Cogn Neurosci Rev 3(2), 71–100. doi: 10.1177/1534582304267187.

Domes, G., Heinrichs, M., Glascher, J., Buchel, C., Braus, D.F., and Herpertz, S.C. (2007a). Oxytocin attenuates amygdala responses to emotional faces regardless of valence. Biological Psychiatry 62(10), 1187–1190. doi: 10.1016/j.biopsych.2007.03.025.

Domes, G., Heinrichs, M., Michel, A., Berger, C., and Herpertz, S.C. (2007b). Oxytocin improves "mind-reading" in humans. Biological Psychiatry 61(6), 731–733.

Domes, G., Hollerbach, P., Vohs, K., Mokros, A., and Habermeyer, E. (2013). Emotional Empathy and Psychopathy in Offenders: An Experimental Study. Journal of Personality Disorders 27(1), 67–84.

Dziobek, I., Rogers, K., Fleck, S., Bahnemann, M., Heekeren, H.R., Wolf, O.T., et al. (2008). Dissociation of cognitive and emotional empathy in adults with asperger syndrome using the multifaceted empathy test (MET). Journal of Autism and Developmental Disorders 38(3), 464–473. doi: 10.1007/s10803-007-0486-x.

Eklund, A., Nichols, T.E., and Knutsson, H. (2016). Cluster failure: Why fMRI inferences for spatial extent have inflated false-positive rates. Proc Natl Acad Sci U S A 113(28), 7900–7905. doi: 10.1073/pnas.1602413113.

Edele, A., Dziobek, I., and Keller, M. (2013). Explaining altruistic sharing in the dictator game: The role of affective empathy, cognitive empathy, and justice sensitivity. Learning and Individual Differences 24, 96–102. doi: 10.1016/j.lindif.2012.12.020.

Fan, Y., Duncan, N.W., de Greck, M., and Northoff, G. (2011). Is there a core neural network in empathy? An fMRI based quantitative meta-analysis. Neuroscience and Biobehavioral Reviews 35(3), 903–911. doi: 10.1016/j.neubiorev.2010.10.009.

Feeser, M., Fan, Y., Weigand, A., Hahn, A., Gärtner, M., Böker, H., et al. (2015). Oxytocin improves mentalizing-pronounced effects for individuals with attenuated ability to empathize. Psychoneuroendocrinology 53, 223–232.

Gamer, M., Zurowski, B., and Buchel, C. (2010). Different amygdala subregions mediate valence-related and attentional effects of oxytocin in humans. Proceedings of the National Academy of Sciences of the United States of America 107(20), 9400–9405. doi: 10.1073/pnas.1000985107.

Gao, S., Becker, B., Luo, L., Geng, Y., Zhao, W., Yin, Y., et al. (2016). Oxytocin, the peptide that bonds the sexes also divides them. Proc Natl Acad Sci US A 113(27), 7650–7654. doi: 10.1073/pnas.1602620113.

Guastella, A.J., Hickie, I.B., McGuinness, M.M., Otis, M., Woods, E.A., Disinger, H.M., et al. (2013). Recommendations for the standardisation of oxytocin nasal administration and guidelines for its reporting in human research. Psychoneuroendocrinology 38(5), 612–625. doi: 10.1016/j.psyneuen.2012.11.019.

Herpertz, S.C., and Bertsch, K. (2014). The social-cognitive basis of personality disorders. Curr Opin Psychiatry 27(1), 73–77. doi: 10.1097/YCO.0000000000000026.

Herpertz, S.C., and Bertsch, K. (2016). Oxytocin Effects on Brain Functioning in Humans. Biological Psychiatry 79(8), 631–632.

Hillis A.E. (2014). Inability to empathize: brain lesions that disrupt sharing and understanding another's emotions. Brain 137, 981–997.

Hu, J.H., Qi, S., Becker, B., Luo, L.Z., Gao, S., Gong Q.Y, et al. (2015). Oxytocin Selectively Facilitates Learning with Social Feedback and Increases Activity and Functional Connectivity in Emotional Memory and Reward Processing Regions. Human Brain Mapping 36(6), 2132–2146.

Hurlemann, R., Patin, A., Onur, O.A., Cohen, M.X., Baumgartner, T., Metzler, S., et al. (2010). Oxytocin Enhances Amygdala-Dependent, Socially Reinforced Learning and Emotional Empathy in Humans. Journal of Neuroscience 30(14), 4999–5007.

Jessica, L.J., Y. Sasaki; Keiko, Ishii; Mizuho, Shinada; Heejung, S. Kim (2016). gene-culture interation influence of culture and oxytocin receptor gene polymorphism on loneliness. Culture and Brain. doi: 10.1007/s40167-

Kawamura, Y., Liu, X., Shimada, T., Otowa, T., Kakiuchi, C., Akiyama, T., et al. (2011). Association between oxytocin receptor gene polymorphisms and autistic traits as measured by the Autism J Spectrum Quotient in a nonJclinical Japanese population. Asia Pacific Psychiatry 3(3), 128–136.

Kim, H.S., Sherman, D.K., Sasaki, J.Y., Xu, J., Chu, T.Q., Ryu, C., et al. (2010). Culture, distress, and oxytocin receptor polymorphism (OXTR) interact to influence emotional support seeking. Proceedings of the National Academy of Sciences of the United States of America 107(36), 15717–15721. doi: 10.1073/pnas.1010830107.

Kirsch, P., Esslinger, C., Chen, Q., Mier, D., Lis, S., Siddhanti, S., et al. (2005). Oxytocin modulates neural circuitry for social cognition and fear in humans. J Neurosci 25(49), 11489–11493. doi: 10.1523/JNEUROSCI.3984-05.2005.

Lamm, C., Decety, J., and Singer, T. (2011). Meta-analytic evidence for common and distinct neural networks associated with directly experienced pain and empathy for pain. Neuroimage 54(3), 2492–2502. doi: 10.1016/j.neuroimage.2010.10.014.

Lamm, C., Nusbaum, H.C., Meltzoff, A.N., and Decety, J. (2007). What Are You Feeling? Using Functional Magnetic Resonance Imaging to Assess the Modulation of Sensory and Affective Responses during Empathy for Pain. PLOS ONE 2(12).

Lee, J., Zaki, J., Harvey, P.O., Ochsner, K., and Green, M.F. (2011). Schizophrenia patients are impaired in empathic accuracy. Psychological Medicine 41(11), 2297–2304.

Leigh, R., Oishi, K., Hsu, J., Lindquist, M., Gottesman, R.F., Jarso, S., et al. (2013). Acute lesions that impair affective empathy. Brain 136, 2539–2549.

Luo, L., Becker, B., Geng, Y., Zhao, Z., Gao, S., Zhao, W., et al. (2017). Sex-dependent neural effect of oxytocin during subliminal processing of negative emotion faces. Neuroimage 162, 127–137. doi: 10.1016/j.neuroimage.2017.08.079.

Luo, S.Y., Li, B.F., Ma, Y., Zhang, W.X., Rao, Y., and Han, S.H. (2015a). Oxytocin receptor gene and racial ingroup bias in empathy-related brain activity. Neuroimage 110, 22–31.

Luo, S.Y., Ma, Y.N., Liu, Y., Li, B.F., Wang, C.B., Shi, Z.H., et al. (2015b). Interaction between oxytocin receptor polymorphism and interdependent culture values on human empathy. Social Cognitive and Affective Neuroscience 10(9), 1273–1281. doi: 10.1093/scan/nsv019.

McLaren, D.G., Ries, M.L., Xu, G., and Johnson, S.C. (2012). A generalized form of context-dependent psychophysiological interactions (gPPI): a comparison to standard approaches. Neuroimage 61(4), 1277–1286. doi: 10.1016/j.neuroimage.2012.03.068.

Melchers, M., Li, M., Chen, Y., Zhang, W., and Montag, C. (2015). Low empathy is associated with problematic use of the Internet: Empirical evidence from China and Germany. Asian journal of psychiatry 17, 56–60.

Montag, C., Sindermann, C., Melchers, M., Jung, S., Luo, R., Becker, B., et al. (2017). A functional polymorphism of the OXTR gene is associated with autistic traits in Caucasian and Asian populations. Am J Med Genet B Neuropsychiatr Genet 174(8), 808–816. doi: 10.1002/ajmg.b.32596.

Panksepp, J., and Panksepp, J.B. (2013). Toward a cross-species understanding of empathy. Trends Neurosci 36(8), 489–496. doi: 10.1016/j.tins.2013.04.009.

Penton-Voak, I.S., Perrett, D.I., Castles, D.L., Kobayashi, T., Burt, D.M., Murray, L.K., et al. (1999). Menstrual cycle alters face preference. Nature 399(6738), 741–742. doi: 10.1038/21557.

Pfundmair, M., Aydin, N., Frey, D., and Echterhoff, G. (2014). The interplay of oxytocin and collectivistic orientation shields against negative effects of ostracism. Journal of Experimental Social Psychology 55, 246–251.

Quintana, D.S., Westlye, L.T., Alnaes, D., Rustan, O.G., Kaufmann, T., Smerud, K.T., et al. (2016). Low dose intranasal oxytocin delivered with Breath Powered device dampens amygdala response to emotional stimuli: A peripheral effect-controlled within-subjects randomized dose-response fMRI trial. Psychoneuroendocrinology 69, 180–188. doi: 10.1016/j.psyneuen.2016.04.010.

Quintana, D.S., Westlye, L.T., Hope, S., Naerland, T., Elvsashagen, T., Dorum, E., et al. (2017). Dose-dependent social-cognitive effects of intranasal oxytocin delivered with novel Breath Powered device in adults with autism spectrum disorder: a randomized placebo-controlled double-blind crossover trial. Transl Psychiatry 7(5), e1136. doi: 10.1038/tp.2017.103.

Radke, S., and de Bruijn, E.R. (2015). Does oxytocin affect mind-reading? A replication study. Psychoneuroendocrinology 60, 75–81. doi: 10.1016/j.psyneuen.2015.06.006.

Reniers, R.L.E.P., Corcoran, R., Drake, R., Shryane, N., and Vollm, B. (2010). The QCAEP: A questionnaire of cognitive and affective empathy. Journal of Personality Assessment 93(1), 84–95.

Riem M.M.E. (2012). Oxytocin Modulates Amygdala, Insula, and Inferior Frontal Gyrus Responses to Infant Crying: A Randomized Controlled Trial (vol 70, pg 291, 2011). Biological Psychiatry 71(7), 660–660. doi: 10.1016/j.biopsych.2012.02.010.

Riem, M.M.E., Bakermans-Kranenburg, M.J., Voorthuis, A., and van IJzendoorn, M.H. (2014). Oxytocin effects on mind-reading are moderated by experiences of maternal love withdrawal: An fMRI study. Progress in Neuro-Psychopharmacology & Biological Psychia try 51, 105–112.

Riem, M.M.E., van IJzendoorn, M.H., Tops, M., Boksem, M.A.S., Rombouts, S.A.R.B., and Bakermans-Kranenburg, M.J. (2012). No Laughing Matter: Intranasal Oxytocin Administration Changes Functional Brain Connectivity during Exposure to Infant Laughter. Neuropsychopharmacology 37(5), 1257–1266. doi: 10.1038/npp.2011.313.

Rilling, J.K., DeMarco, A.C., Hackett, P.D., Thompson, R., Ditzen, B., Patel, R., et al. (2012). Effects of intranasal oxytocin and vasopressin on cooperative behavior and associated brain activity in men. Psychoneuroendocrinology 37(4), 447–461. doi: 10.1016/j.psyneuen.2011.07.013.

Rodrigues, S.M., Saslow, L.R., Garcia, N., John, O.P., and Keltner, D. (2009). Oxytocin receptor genetic variation relates to empathy and stress reactivity in humans. Proceedings of the National Academy of Sciences of the United States of America 106(50), 21437–21441. doi: 10.1073/pnas.0909579106.

Rosenfeld, A.J., Lieberman, J.A., and Jarskog, L.F. (2011). Oxytocin, Dopamine, and the Amygdala: A Neurofunctional Model of Social Cognitive Deficits in Schizophrenia. Schizophrenia Bulletin 37(5), 1077–1087. doi: 10.1093/schbul/sbq015.

Scheele, D., Kendrick, K.M., Khouri, C., Kretzer, E., Schlapfer, T.E., Stoffel-Wagner, B., et al. (2014). An oxytocin-induced facilitation of neural and emotional responses to social touch correlates inversely with autism traits. Neuropsychopharmacology 39(9), 2078–2085. doi: 10.1038/npp.2014.78.

Schulte-Ruther, M., Markowitsch, H.J., Fink, G.R., and Piefke, M. (2007). Mirror neuron and theory of mind mechanisms involved in face-to-face interactions: A functional magnetic resonance imaging approach to empathy. Journal of Cognitive Neuroscience 19(8), 1354–1372. doi: DOI 10.1162/jocn.2007.19.8.1354.

Shamay-Tsoory S.G. (2011). The neural bases for empathy. Neuroscientist 17(1), 18–24. doi: 10.1177/1073858410379268.

Shamay-Tsoory, S.G., and Abu-Akel, A. (2016). The Social Salience Hypothesis of Oxytocin. Biological Psychiatry 79(3), 194–202.

Shamay-Tsoory, S.G., Abu-Akel, A., Palgi, S., Sulieman, R., Fischer-Shofty, M., Levkovitz, Y., et al. (2013). Giving peace a chance: Oxytocin increases empathy to pain in the context of the Israeli-Palestinian conflict. Psychoneuroendocrinology 38(12), 3139–3144. doi: 10.1016/j.psyneuen.2013.09.015.

Shamay-Tsoory, S.G., Aharon-Peretz, J., and Levkovitz, Y. (2007). The neuroanatomical basis of affective mentalizing in schizophrenia: Comparison of patients with schizophrenia and patients with localized prefrontal lesions. Schizophrenia Research 90(1-3), 274–283. doi: 10.1016/j.schres.2006.09.020.

Shamay-Tsoory, S.G., Aharon-Peretz, J., and Perry, D. (2009). Two systems for empathy: a double dissociation between emotional and cognitive empathy in inferior frontal gyrus versus ventromedial prefrontal lesions. Brain 132(Pt 3), 617–627. doi: 10.1093/brain/awn279.

Singer, T., and Lamm, C. (2009). The Social Neuroscience of Empathy. Year in Cognitive Neuroscience 2009 1156, 81–96.

Singer, T., Snozzi, R., Bird, G., Petrovic, P., Silani, G., Heinrichs, M., et al. (2008). Effects of Oxytocin and Prosocial Behavior on Brain Responses to Direct and Vicariously Experienced Pain. Emotion 8(6), 781–791. doi: 10.1037/a0014195.

Slotnick S.D. (2017). Cluster success: fMRI inferences for spatial extent have acceptable false-positive rates. Cogn Neurosci 8(3), 150–155. doi: 10.1080/17588928.2017.1319350.

Smith, K.E., Porges, E.C., Norman, G.J., Connelly, J.J., and Decety, J. (2014). Oxytocin receptor gene variation predicts empathic concern and autonomic arousal while perceiving harm to others. Social Neuroscience 9(1), 1–9. doi: 10.1080/17470919.2013.863223.

Sobota, R., Mihara, T., Forrest, A., Featherstone, R.E., and Siegel, S.J. (2015). Oxytocin reduces amygdala activity, increases social interactions, and reduces anxiety-like behavior irrespective of NMDAR antagonism. Behav Neurosci 129(4), 389–398. doi: 10.1037/bne0000074.

Spengler, F.B., Schultz, J., Scheele, D., Essel, M., Maier, W., Heinrichs, M., et al. (2017). Kinetics and Dose Dependency of Intranasal Oxytocin Effects on Amygdala Reactivity. Biol Psychiatry 82(12), 885–894. doi: 10.1016/j.biopsych.2017.04.015.

Spielberger, C.D., Gorsuch, R.L., and Lushene, R.E. (1970). Manual for the state-trait anxiety inventory.

Stern, R.M., Ray, W.J., and Quigley, K.S. (2001). Psychophysiological recording. Oxford University Press, USA.

Striepens, N., Kendrick, K.M., Maier, W., and Hurlemann, R. (2011). Prosocial effects of oxytocin and clinical evidence for its therapeutic potential. Frontiers in Neuroendocrinology 32(4), 426–450. doi: 10.1016/j.yfrne.2011.07.001.

Striepens, N., Scheele, D., Kendrick, K.M., Becker, B., Schafer, L., Schwalba, K., et al. (2012). Oxytocin facilitates protective responses to aversive social stimuli in males. Proc Natl Acad Sci U S A 109(44), 18144–18149. doi: 10.1073/pnas.1208852109.

Tully, J., Gabay, A.S., Brown, D., Murphy, D.G.M., and Blackwood, N. (2018). The effect of intranasal oxytocin on neural response to facial emotions in healthy adults as measured by functional MRI: A systematic review. Psychiatry Res 272, 17–29. doi: 10.1016/j.pscychresns.2017.11.017.

Uddin L.Q. (2015). Salience processing and insular cortical function and dysfunction. Nat Rev Neurosci 16(1), 55–61. doi: 10.1038/nrn3857.

Watson, D., Clark, L.A., and Tellegen, A. (1988). Development and validation of brief measures of positive and negative affect: the PANAS scales. J Pers Soc Psychol 54(6), 1063–1070.

Weisman, O., Pelphrey, K.A., Leckman, J.F., Feldman, R., Lu Y, Chong, A., et al. (2015). The association between 2D:4D ratio and cognitive empathy is contingent on a common polymorphism in the oxytocin receptor gene (OXTR rs53576). Psychoneuroendocrinology 58, 23–32. doi: 10.1016/j.psyneuen.2015.04.007.

Wigton, R., Radua, J., Allen, P., Averbeck, B., Meyer-Lindenberg, A., McGuire, P., et al. (2015). Neurophysiological effects of acute oxytocin administration: systematic review and meta-analysis of placebo-controlled imaging studies. Journal of Psychiatry & Neuroscience 40(1), E1–E22.

Wingenfeld, K., Kuehl, L.K., Janke, K., Hinkelmann, K., Dziobek, I., Fleischer, J., et al. (2014). Enhanced Emotional Empathy after Mineralocorticoid Receptor Stimulation in Women with Borderline Personality Disorder and Healthy Women. Neuropsychopharmacology 39(8), 1799–1804. doi: 10.1038/npp.2014.36.

Wong, C.-S., and Law, K.S. (2002). The effects of leader and follower emotional intelligence on performance and attitude: An exploratory study. The leadership quarterly 13(3), 243–274.

Wolf, O.T., Schulte, J.M., Drimalla, H., Hamacher-Dang, T.C., Knoch, D., and Dziobek, I. (2015). Enhanced emotional empathy after psychosocial stress in young healthy men. Stress 18(6), 631–637. doi: 10.3109/10253890.2015.1078787.

Wu, N., Li, Z., and Su, YJ. (2012). The association between oxytocin receptor gene polymorphism (OXTR) and trait empathy. Journal of Affective Disorders 138(3), 468–472. doi: 10.1016/j.jad.2012.01.009.

Xu, L., Ma, X., Zhao, W., Luo, L., Yao, S., and Kendrick, K.M. (2015). Oxytocin enhances attentional bias for neutral and positive expression faces in individuals with higher autistic traits. Psychoneuroendocrinology 62, 352–358. doi: 10.1016/j.psyneuen.2015.09.002.

Xu, X., Yao, S., Xu, L., Geng, Y., Zhao, W., Ma, X., et al. (2017). Oxytocin biases men but not women to restore social connections with individuals who socially exclude them. Scientific reports 7, 40589.

Young, L.J., and Barrett, C.E. (2015). Neuroscience. Can oxytocin treat autism? Science 347(6224), 825–826. doi: 10.1126/science.aaa8120.

Zhao, W., Geng, Y., Luo, L., Zhao, Z., Ma, X., Xu, L., et al. (2017). Oxytocin Increases the Perceived Value of Both Self-and Other-Owned Items and Alters Medial Prefrontal Cortex Activity in an Endowment Task. Front Hum Neurosci 11, 272. doi: 10.3389/fnhum.2017.00272.

